# Evolution and developmental diversity of skin spines in pufferfish

**DOI:** 10.1101/347690

**Authors:** Takanori Shono, Alexandre P. Thiery, Daisuke Kurokawa, Ralf Britz, Gareth J. Fraser

## Abstract

Teleost fishes develop a huge variety of skin ornaments. How these diverse skin structures develop in fishes is unknown. The teleost fish order Tetraodontiformes includes some of the most unusual fishes such as the ocean sunfish, triggerfish and pufferfish, and they all can develop a vast assortment of scale derivatives that cover their bodies. Pufferfish have some of the most extreme scale derivatives, dermal spines, which are erected during their characteristic puffing behavior. Here we show that pufferfish spines develop through conserved gene interactions essential for other vertebrate skin appendage formation, like hair and feathers. However, pufferfish spines form without EDA (ectodysplasin), an essential molecule for the development of most vertebrate skin appendages. Modifying signaling pathways lead to loss or reduction of spine coverage in pufferfish, suggesting a mechanism for skin appendage diversification. We suggest that pufferfish skin spines evolved from a basic teleost scale-type through derived gene network modification in Tetraodontiformes.

## Introduction

Vertebrates show a great diversity and elaboration of skin ornament and appendages, ranging from the various types of mineralized scales in fishes and keratinous scales of reptiles, to complex feathers in birds and hair in mammals. Despite structural non-homology of most of these units and the apparent uniqueness to each vertebrate clade*^1–3^*, components of the genetic control of skin appendage patterning and development appear to be conserved and shared among these diverse vertebrate clades*^4^*. The development of skin appendages such as hair follicles, teeth, mammary glands, feathers and epidermal scales in tetrapod lineages have been well investigated*^4–7^* and these studies imply that the respective developmental process and molecular mechanisms are deeply conserved among vertebrate skin appendages. The earliest developmental stages of vertebrate skin appendages, generally, are remarkably similar in their initial morphogenesis of the initial placode, which is then followed by specific divergence of an assortment of epithelial units*^7^*. Morphogenesis of these units depends on inductive epithelial-mesenchymal interactions mediated by a conserved set of signaling molecules, associated with the Hedgehog (Hh), Wnt, Bmp, FGF, Notch and EDA pathways*^7^*. Skin appendages in tetrapod lineages arise from epithelial cells of ectodermal origin*^4^* whereas in contrast teleost scales are produced initially from underlying mesodermal layers of the dermis *^6, 8, 9^*, however the true extent of teleost scale initiation is currently unknown.

Although the diversity of skin ornaments in teleost fishes is vast, we currently know little about the development of these structures that include a diverse array of scales, and scale-like skin appendages*^10^*. The molecular development of dermal scales of teleost fishes is surprisingly under-studied despite their diversity and superficial formation on the surface of the body. The potential reason for this lack of comparative knowledge could be the relatively late developmental timing of scale formation during early juvenile stages in most teleosts, including the zebrafish. However some signaling pathways have been documented*^11–15^*. For instance, during scale development in zebrafish (*Danio rerio*) the ligand of the Hh pathway *shh* is expressed in the epidermal layer of cells adjacent to the scale primordium in the mesoderm*^10^*. More recently a set of studies documenting the association of the Wnt/β-catenin and the Fgf signaling pathway in teleost scale development and morphogenesis have emerged^16–18^. In addition, mutant phenotypes of the *edar* (Ectodyplastin-A receptor) coding locus in Medaka (*Oryzias latipes*; Beloniformes) show an almost complete loss of scale formation suggesting that the eda/edar signaling pathway is necessary for normal scale formation in teleosts*^12^*. Eda/edar signaling also controls morphology and patterning of the body plate armour in both marine and freshwater populations of the three-spined stickleback*^14,19^*. Further information on the developmental significance of Ectodysplasin pathway members (*eda* and *edar*) on the patterning of adult scales in zebrafish has recently been documented*^11^*, where eda/edar have been shown to be important in regulating signaling centers from the organization of epidermal cells that are responsible for scale development. The involvement of these common genetic programs during the development of epidermal/dermal skin appendages from diverse clades of vertebrates suggests molecular similarity of skin patterning mechanisms. As a corollary, we expect that even the most extremely modified skin ornament in teleosts shares elements of the core developmental skin patterning programs with that of other vertebrate skin appendages.

Tetraodontiformes include some of the most unusual-looking teleost fishes such as ocean sunfishes (Molidae), boxfishes (Ostraciidae) and porcupine fishes (Diodontidae). With approximately 350 species^*20, 21*^, Tetraodontiformes epitomize extreme morphological diversification, which includes, but is not limited to, the craniofacial and dermal skeleton*^21, 22^* Members of each of the tetraodontiform clades have a highly unusual covering of their integument by mineralized structures ranging from spinoid scales to individual spines and even thick, plate-like armor (Fig. 1). Dermal scale development in Tetraodontiformes is thought to be associated with evolution and modification of the typical squamous scale types (e.g. cycloid, spinoid or ctenoid) found in most of teleost fishes*^23^*, however evolutionary developmental approaches have not been used so far to test this hypothesis.

**Fig. 1.**
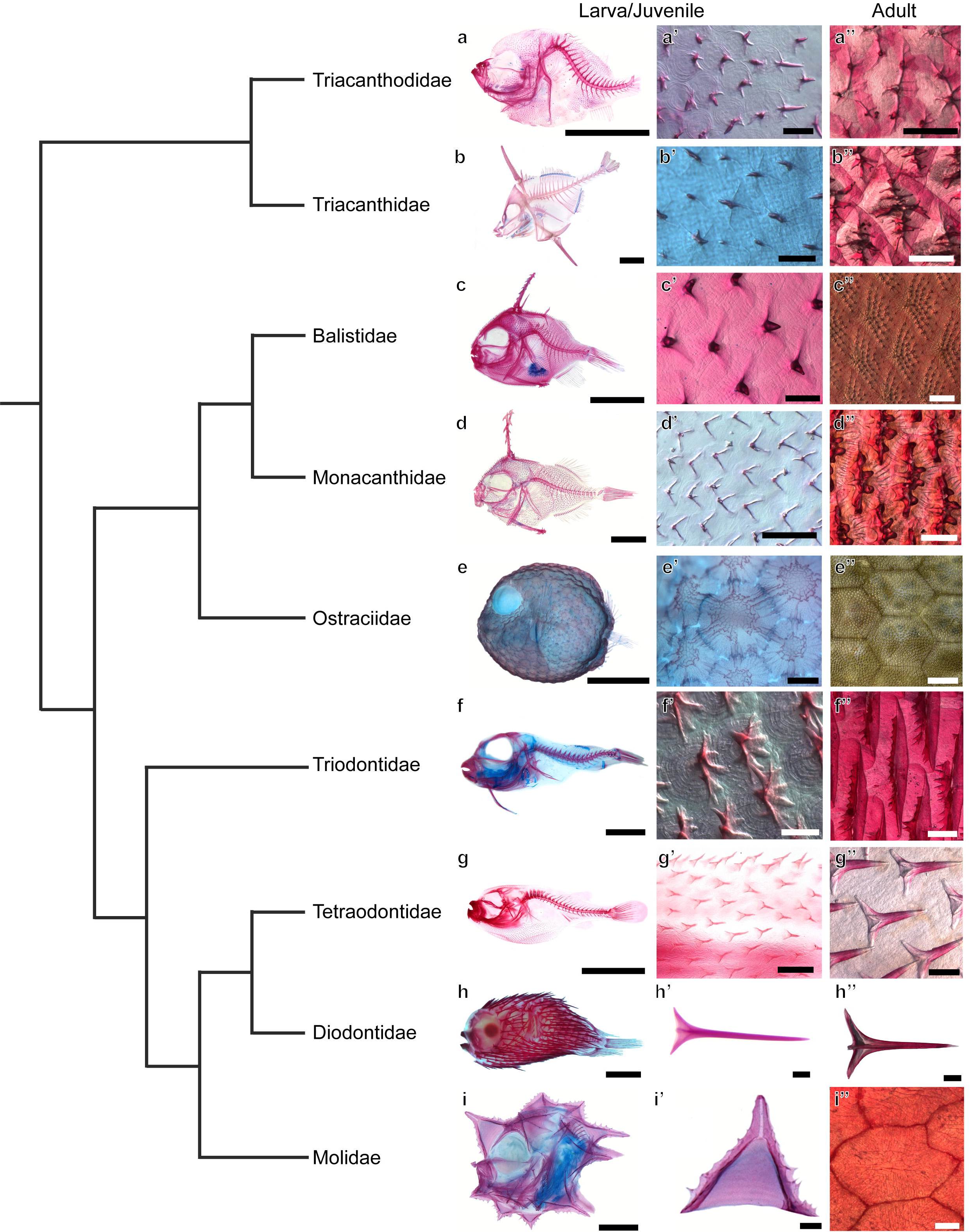
Skin appendage diversity in the Order Tetraodontiformes. Cleared and stained spinoid scales and spines of larvae/juveniles and adults in representative species of tetraodontiform families. a Triacanthodidae, *Hollardia* sp., Standard Length (SL); 5mm (**a, a’**) and *Paratriacanthodes herrei*, SL; 42.5 mm (**a”**). **b** Triacanthidae, *Tripodichthys oxycephalus*, SL; 4, 26 mm (**b, b’**) and *Pseudotriacanthus strigilifer* SL; 88 mm (**b”**). **c** Balistidae, *Balistes vetula*, SL; 10 mm (**c, c’**) and *B. capriscus*, SL; 128 mm (**c”**). **d** Monacanthidae, *Stephanolepis* sp., SL; 5 mm (**d, d’**) and *S. hispidus* SL; 94.5 mm. **e** Ostraciidae, *Ostracion* sp., SL; 6.2 mm (**e, e’**) and *Ostracion trigonus*, SL; 330 mm (**e”**). **f** Triodontidae, *Triodon macropterus*, SL; 20 mm (**f, f’**) and *T. macropterus* SL; 315 mm (**f”**). g Tetraodontidae, *Takifugu niphobles*., 10 mm (**g, g’**) and *T. niphobles*, SL; 102.3 mm (**g”**). **h** Diodontidae, *Diodon holocanthus*, SL; 12.5 mm (**h, h’**) and *D. holacanthus* SL; 101 mm (**h”**). **i** Molidae, *Ranzania laevis*, SL; 1.7 mm (**i, i’**) and *R. laevis*, SL; 620 mm (**i”**). Phylogenetic relationships of tetraodontiform families follow a published data^*17*^. Scale bars, 2mm (**a, b, d, e, g, h**), 500 μ.m (**a’, a”, b”, g”, i**) 5mm (**b, f**) 100 μm (**b’, c’, d’, f’, g’, h’, i’**), 1 mm (**c”, e”, f”, h”, i”**), 200 μm (**d”, e’**).

The modification of scale derivatives in Tetraodontiformes has produced extreme forms, and porcupine fishes (Diodontidaes), for example, are aptly named for their conspicuous skin appendages. Erection of their dermal spines (Fig. 1) is linked to their ability to inflate, producing a powerful defense mechanism^*24*^. This mechanism for inflation is shared with the pufferfishes (Tetraodontidae), a group that shows much less impressive but more diverse body spination^*21*^. To further our understanding of the development of dermal skin coverings in tetraodontids, we focus on one species of pufferfish, *Takifugu niphobles*. We investigate the formation of its body spination, which is restricted to the ventral body surface in the early juvenile stage and reinforces the abdominal region during inflation. The *Takifugu* pufferfish lineage is well known from recent genome sequencing efforts which demonstrated that (i) pufferfishes represent the vertebrate with the smallest, most compact genome, and (ii) that the *Takifugu* lineage has undergone a relatively recent explosive radiation *^25,26^*. Even within the pufferfish clade there is a variation in spine size and shape, as well as in the level of spine coverage across the body, ranging from a complete lack of spines to a dense, full body coverage^*21*^.

Here we present (i) information on the diversity and modification of skin ornament in this particularly diverse order of teleost fishes, the Tetraodontifomes; describe (ii) the development of the dermal spines in the derived Tetraodontiform family, pufferfish (Tetraodontidae; *Takifugu niphobles*) as a model for skin appendage novelty and diversification; and report (iii) on the results of the developmental and genetic manipulation of the spines that cover the ventral surface of *Takifugu* and finally (iv) suggest a mechanism for how morphological diversity of skin ornament in this group may have arisen.

## Results

### Diverse Morphological Structure of Skin Appendages in the Order Tetraodontiformes

The morphology of dermal ornament, i.e. spines, plates and scales, and their localization on the body surface vary greatly between members of the 10 families of the order Tetraodontiformes; Triacanthodidae (spikefishes), Triacanthidae (triple spine fishes or tripodfishes), Balistidae (triggerfishes), Monacanthidae (filefishes), Aracanidae (boxfishes), Ostraciidae (true boxfishes), Triodontidae (three-tooth pufferfishes), Tetraodontidae (four-tooth, true pufferfishes), Diodontidae (porcupine fishes) and Molidae (ocean sunfishes)^*20,27*^. To understand spine or scale structure in the order Tetraodontiformes, we studied cleared and double-stained larval/juvenile and adult specimens, from species representing the all 9 of 10 extant ttetraodontiform families (Fig. 1). The morphology and localization of dermal structures in juveniles were similar to that in adults in Triodontidae, Tetraodontidae and Diodontidae, but this is not the case in other groups. The juveniles of both the basal spikefishes (family Triacanthodidae) and the more derived three-toothed pufferfish (family Triodontidae) exhibit mineralized units on the body surface that consist of a scale like plate with one (in triacanthodids) or several (in triodontids) individual spines at the posterior end. This plate strongly resembles a typical cycloid scale in having concentric ridges, the so-called circuli (Fig. 1a, f). This “cycloid” base in these two groups is still present, though less obvious, in the spinoid scales observed in the adults^*28*^. Juveniles of the tripod fishes (family Triacanthidae), the triggerfishes (family Balistidae), and the filefishes (family Monacanthidae) also exhibit a single spined unit on a “cycloid” base, which, however, appear to lack circuli. In contrast, in adults of members of these three families a more complex structure is present, with spines or spinules sometimes covering the entire visible scale surface (Fig. 1c, d). Boxfishes (family Ostraciidae) do not possess any spines, but instead have a covering of separate hexagonal bony plates on the body surface of juveniles, which grow together to form a thick armor in adults (Fig. 1e) *^21^*. Pufferfishes (family Tetraodontidae) and porcupine fishes (family Diodontidae) possess individual spines with their spatial position and arrangement varying among species. While porcupine fishes have particularly long spines on their entire body, those of pufferfishes range from complete coverage over the entire body (seen in *Carinotetraodon^29^*), to a restriction of spines to certain regions of the body. In *Takifugu*, these spines are restricted to the dorsal head and ventral abdominal regions. Individual spines of pufferfishes are composed of a basal multi-pronged root embedded into the dermis and a distally pointed single spine tip (Fig. 1g, h). Ocean sunfish (family Molidae) juveniles are covered with a network of jagged edged pyramidal spines covering the skin, whereas adults have no spines, but instead their entire body is covered with polygonal scale-like plates (Fig. 1i) *^30^*. In summary, Tetraodontiformes show a diversity in dermal skin ossifications that rivals that of many other, more species-rich teleost orders (see Supplementary Table 1 for a summary of observations) and for this reason are especially suited for the study of evolutionary and developmental diversification of body ornamentation.

### Pufferfish spines develop from early mesenchymal primordia

Morphological descriptions of spinoid scales, plates and spines have previously been reported predominantly for adult Tetraodontiformes ^*20,21,23,28,31–37*^. However, how this variety of dermal appendages develop remains to be studied with a specific developmental approach. Therefore, to begin this process we investigated the histology and development of spines using one species of pufferfish, the Japanese Grass Pufferfish (*Takifugu niphobles*, family: Tetraodontidae). Scanning electron microscopy (SEM) revealed a bumpy skin surface in embryos 12 days post-fertilization (dpf), located between the ventral pectoral girdle and the abdominal region (Fig. 2a). A close-up of this region highlights ectodermal units that project from the body surface marking the initial site of development of a spine primordium at this stage (Fig. 2b). Sagittal sections of embryos at 13 dpf demonstrated that the developing spine regions are composed of two tissue layers, epithelium and mesenchyme, and associated pigment cells e.g melanocytes. The spine primordia derive from mesenchyme (dermis) and are strongly stained with haematoxylin (and Alcian Blue; Fig. 2c). In juvenile stages (at 46 dpf), developing spines extend from a fibrous layer of the dermis (Fig. 2d). In adult stages, Alizarin Red-positive, well-mineralized, functional spines extend to and protrude through the outer layers of the skin (Fig. 2e). These observations, coupled with our results from tissue staining assays show that spines develop below the epidermis in *Takifugu* from the mesenchyme (dermal mesoderm) during early ontogeny. This is similar to the development of regular teleost scales. In zebrafish (*Danio rerio*; Cypriniformes) and Medaka (*Oryzias latipes*; Beloniformes), scales also develop from dermal mesoderm but do so much later, approximately one-month post-fertilization^*10, 12, 38*^. However, the location of the initial signals for spine/scale competence is debatable, and could arise from the either mesodermal or epidermal origins.

**Fig. 2.**
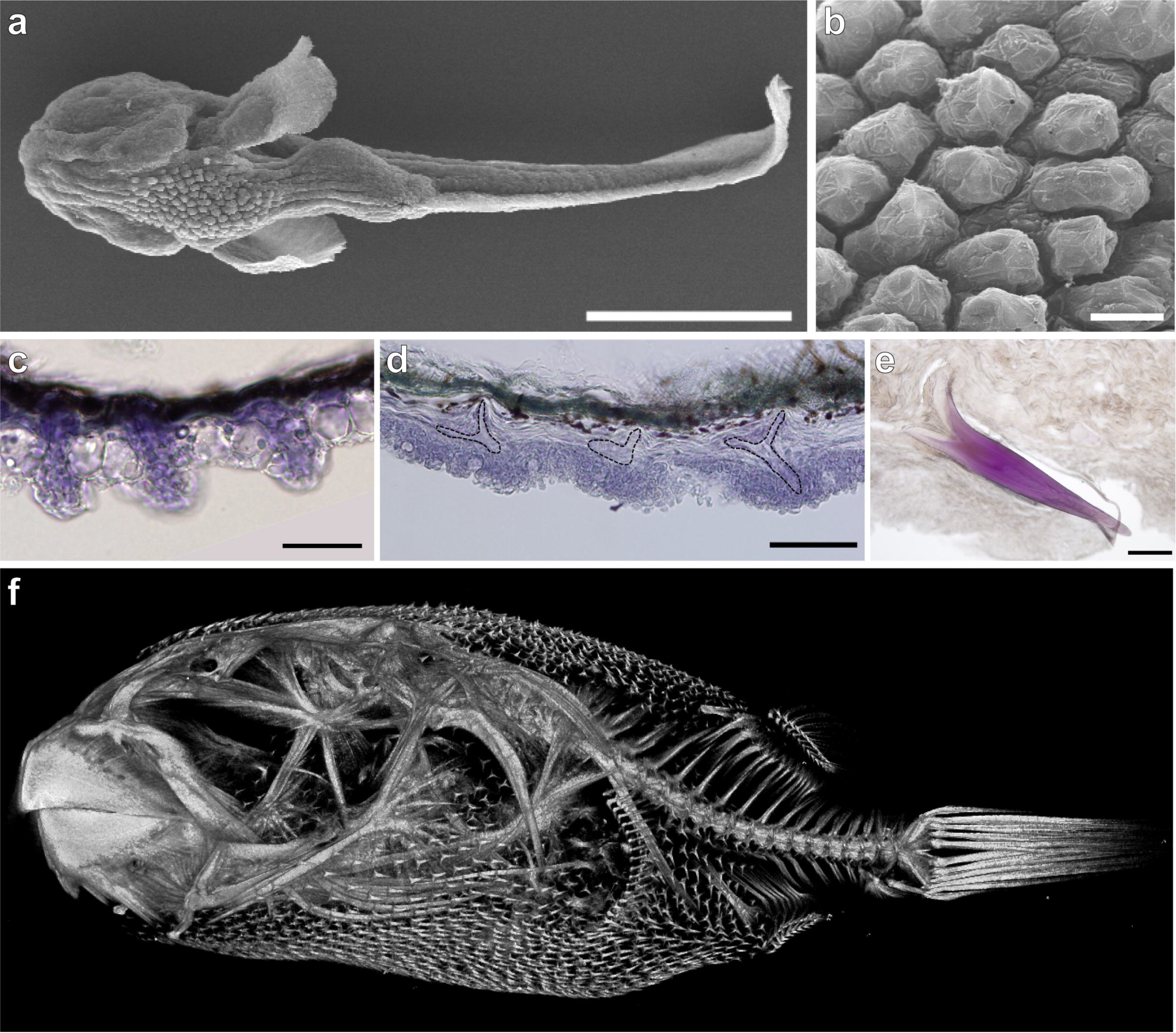
Location and histological structure of spines of *Takifugu*. **a** SEM of ventral bossy surface of *Takifugu niphobles*, 12 dpf. **b** Close-up of spine regions. **c** Sagittal sections of spine regions in 13 dpf of an embryo stained with Haematoxylin and Alcian Blue staining. **d** Sagittal section of spines in 46 dpf embryo stained with Haematoxylin. Black dotted lines indicate spines. **e** Sagittal section of spines in an adult stained with Alizarin Red. **f** DRISHTI rendered MicroCT scan of the adult *Takifugu oblongus*, showing the location of spine coverage on the ventral and dorsal surface. Scale bars, 500 μm (**a**), 20 μm (**b**), 50 μm (**c, d**), 200 μm (**e**).

### Wnt, Hh and FGF pathways are important for spine development

Our histological and developmental observation of spines in larval *Takifugu* suggests that the main component of spine development is mesodermal, similar to the cellular source of scale development in other teleost fishes^*8, 9*^. Therefore, we hypothesize that there is a high level of conservation between spine and scale development in these teleost groups. We speculate that spines form through similar interactions between the epithelium and mesenchyme with genetic pathways equivalent to those during general teleost scale development. This suggests that these distinct sets of scale-type structures are identical in tissue derivation and homologous, despite their morphological divergence.

Because little is known regarding gene expression during scale developmental, we examined a set of gene markers during spine formation in *Takifugu*. To do this we selected candidate genes from major developmental pathways (Hh, Wnt, FGF and BMP signaling pathways) that are known to be involved during organogenesis in a range of structures, i.e. jaws, teeth, limbs, bone and skin (ectodermal) appendages in vertebrates^*7,39–41*^. *β-catenin* is a signal transducer of canonical Wnt signaling, a pathway known to play important roles in cell proliferation and differentiation during organogenesis of various structures^*42*^. We observe expression of *β-catenin* within the spine primordium and in the surrounding epithelial cells (Fig. 3a). The expression of *β-catenin* in spine primordia is stronger at the boundary region between spines and the overlying epithelium. The transcription factor, *lef1* (Lymphoid enhancer binding factor-1), is also associated with the canonical Wnt pathway by interaction with *β-catenin^43^* and is expressed during spine primordium development, broadly in the apical region of the epithelium (Fig. 3b). The spatial expression of *lef1* is consistent throughout the early regional specification that demarcates spine competent ectoderm, with its expression upregulated at the border of the ventrally restricted spine-forming region (Fig. 3b). The *Wnt* pathway ligand, *wnt7a*, of the frizzled family of transmembrane receptors^*44*^, is expressed throughout the both mesenchymal and epithelial compartment, although its expression is stronger at the apical tip of the epithelial where spine forms (Fig. 3c). *shh*, a ligand of the Hh pathway, has a critical role as a morphogen in cell division, specification and patterning of organs^*45*^. *shh* is expressed in the epithelium adjacent to the spine primordia (Fig. 3d). This expression pattern shows similarity with scale development in zebrafish, where *shh* is expressed in the epithelium adjacent to the scale anlagen^*10*^ *Sostdc1* (Sclerostin domain-containing 1, also known as *Wise, ectodin* and *USAG-1*), encodes an N-glycosylated, secreted protein associate with the BMP pathway as an antagonist, and mediates regulation of the network for both the Hh and Wnt pathway^*46, 47*^. Expression of *Sostdc1* is present in the ventral spine region and localization is restricted to the distal end of the spine unit (Fig. 3e). FGF ligands play a key role in the processes of proliferation and differentiation in various organs, interacting with Hh, Wnt and BMP signaling pathways^*48*^. *Fgf3* is expressed in epithelial cells adjacent to the apical tip of the developing spine in *Takifugu* (Fig. 3f). In contrast, *Fgf10a* is expressed extensively in the epithelium, however its expression is also upregulated in the epithelium adjacent to the apical tip of spines (Fig. 3g). The BMP pathway is involved in a variety of cellular interactions during various organogenic processes^*49*^. The BMP ligands *bmp2* and *bmp4* are both expressed in the developing spine and importantly in a pre-patterned ventral restriction prior to spine primordia formation, demarcating the homogenous, yet restricted, field where spines will form (Fig. 3h, i). This *bmp2/4*-marked pre-pattern is intriguing as the formation of spines is restricted to within this region, potentially suggesting that BMP signaling is important in initiating a spine competent dermis in *Takifugu*. Further BMP-associated gene expression shows a restricted expression pattern e.g. *Fst* (*Follistatin*), an activin-binding protein known to be a BMP antagonist^*50-51*^, is expressed in the mesenchyme surrounding each spine primordia (Fig. 3j). These results highlight that common markers of ectodermal appendage formation are expressed and active during initiation, development and organization of the highly derived spine ornament of pufferfishes.

**Fig. 3.**
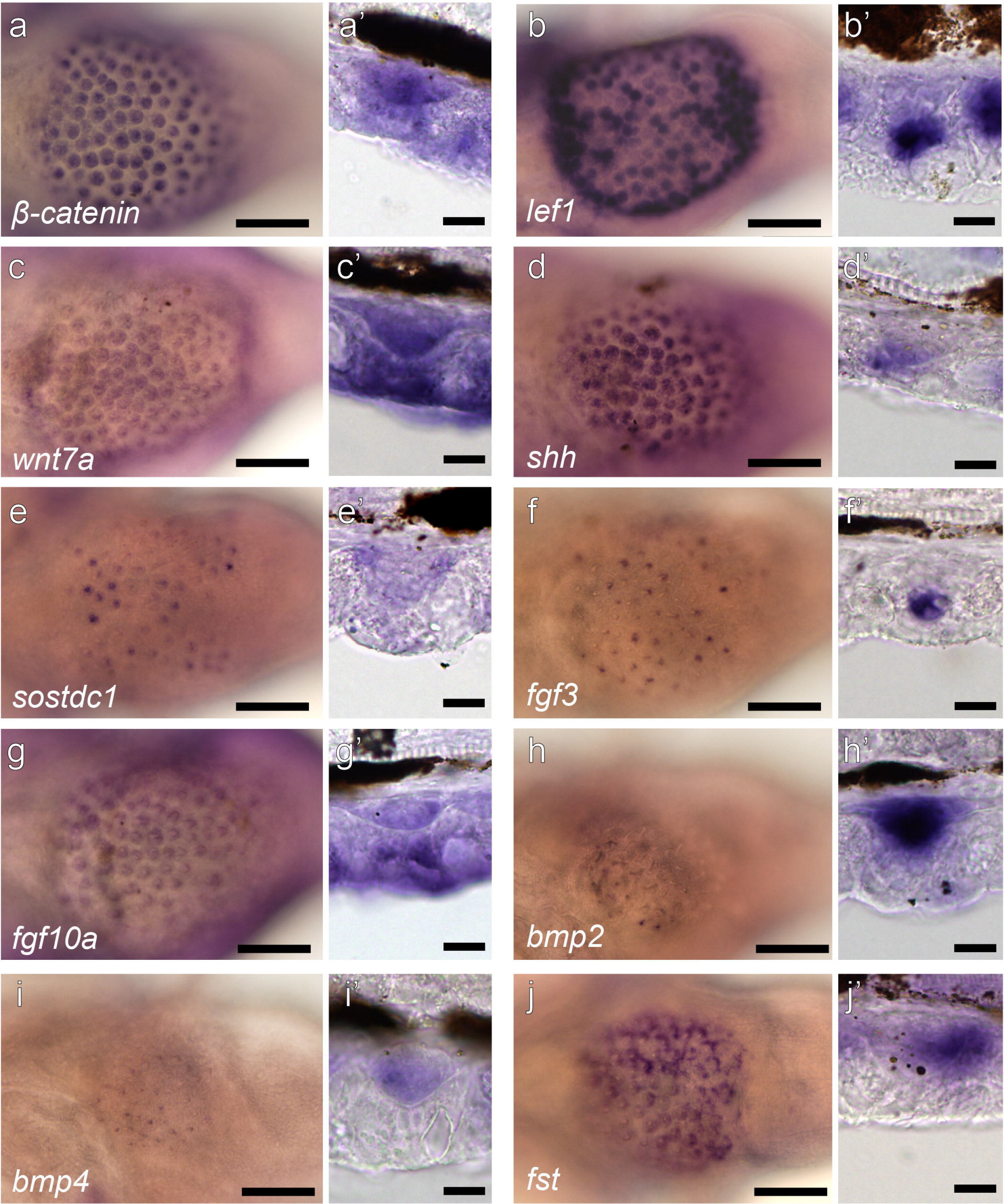
Gene expression in embryonic spine primordia of *Takifugu* at 12 dpf. Gene expression for *β-catenin* (**a**), *lefl* (**b**), *wnt7a* (**c**), *shh* (**d**), *sostdcl* (**e**), *fgf3*(**f**), *fgf10a* (**g**), *bmp2*(**h**), *bmp4*(**i**) and *fst* (**j**). Right above each panel (**a-h**) shows images of spine primordia in sagittal section. White dotted line indicates a boundary line between spine primordium and epithelium layer. Scale bar, 100 μm.

### Functional manipulation of the BMP pathway during Takifugu spine development

Because BMP signals regulate number, pattern and size during the development of skin appendages^*52–54*^, and given the known conservation of genetic markers of ectodermal patterning, we hypothesize that similar genetic mechanisms might regulate dermal spine patterning and development in *Takifugu*. At 10 dpf, the stage before initiation of spine primordia, *bmp2* and *bmp4* are expressed in the ventral mesenchyme of the embryo defining the specific ventral territory (restricted region) where spines form (Fig. 4a, b). This expression pattern suggests that BMP pathway molecules are involved in the initiation or determination of spine patterning. This expression in a ventral patch appears to demarcate a pre-pattern for future spine formation. To identify the functional role of the BMP pathway in spine development, we attempted Bmp gene knock-down assays through the injection of an antisense morpholino (MO) in fertilized *Takifugu* eggs at the one-cell stage. Although injections with *bmp2* or *bmp4-MO* resulted in mortality at the hatching stage (6 dpf), morphants injected with *follistatin (fst)*-MO, an antagonist of Bmp, continued to develop. *Fst*-MO morphants resulted in the formation of abnormal lower jaws at 12 dpf (Fig. 5a, b). Alcian Blue staining revealed that morphants injected with fst-MO developed an abnormal arrangement of Meckel’s cartilage and branchial cartilages with the formation of only four branchial arches, rather than the normal set of five (Fig. 5c, d). In contrast, wild type embryos developed normal cartilages and the standard set of five ceratobranchials. Similarly, the knock down of *fst* via MO injection in zebrafish resulted in a reduction in the number of branchial arches cartilages^*55*^. These data confirm that the *fst* morpholino experiments in pufferfish resulted in known skeletal morphotypes previously observed in *fst* knock down experiments in other teleost fishes^*55*^. However, beyond these known phenotypes we also observed novel phenotypes in *Takifugu* in relation to dermal spine formation. In wild type *Takifugu* embryos, spines are present in the ventral region (Fig. 5e, g) whereas in morphant embryos injected with *fst*-MO, hyperplasia of spines was observed. Furthermore, the apical tip of spines developed earlier than that of the control embryos (Fig. 5f, h). We also documented a reduction in the number of spines in the region (per 2 mm^2^) in morphant with *fst*-MO, when compared with the number of spines in wild type (Numbers are described in Fig. 5i). These results suggest that whilst the spatial patterning of spines in the ventral region was expanded in *Takifugu* morphants injected with *fst*-MO, the overall number of individual spines was reduced (Fig. 5i).

**Fig. 4.**
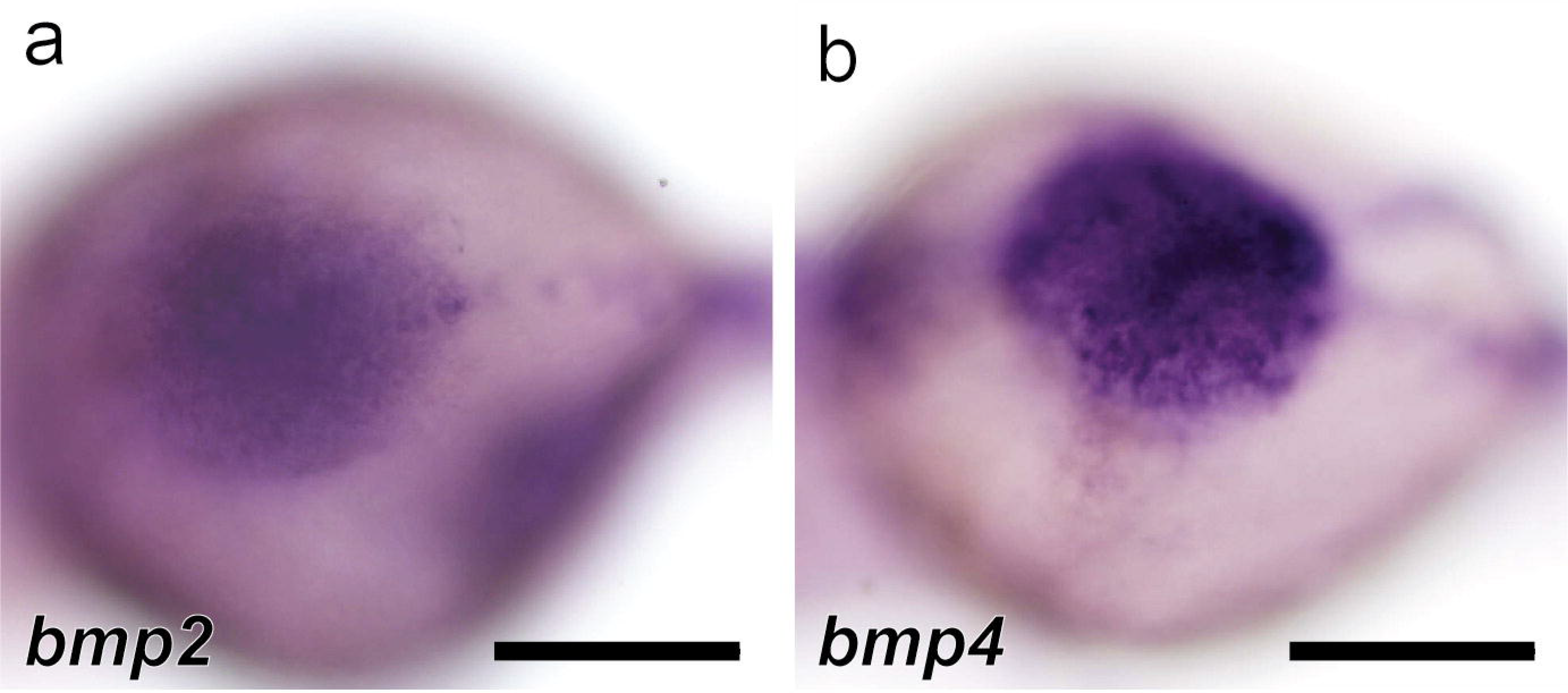
BMP signaling genes demarcate spine competent region in *Takifugu* during early development. Ventral view of 10 dpf embryos. During initiation of spine development, *bmp2* (**a**) and *bmp4* (**b**) are expressed in a specific region of ventral (abdominal) mesenchyme. Scale bar, 200 μm.

**Fig. 5.**
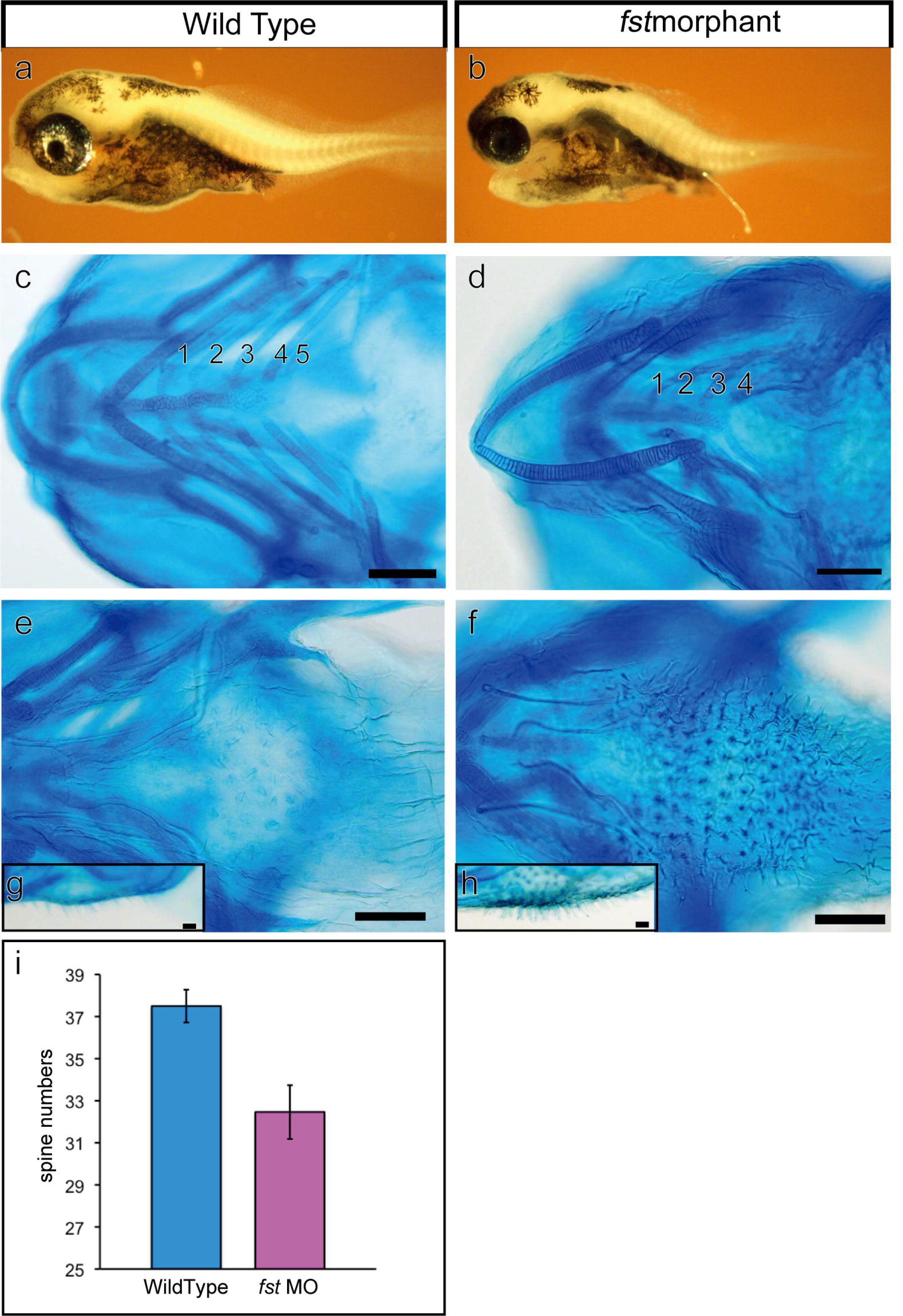
Hypertrophy of spine formation in *Takifugu* embryos after injection of *fst* morpholino. Wild type embryo (**a, c, e, g**) and a morphant with *fst* morpholino (*fst* MO) (**b, d, f, h**). **a-b** Lateral view of embryos. **c-h** Embryos stained with Alcian Blue. **c** Dorsal view of craniofacial skeleton in wild type embryos showing the normal set of five branchial arches, (100%, n = 14). **d** Severe phenotype of a morphant showing an abnormal arrangement of Meckel’s cartilage and only four branchial arches (18.2 %, n = 11). (**e, f**) Ventral view of embryos in the region of spine development. (**g, h**) Close up of spines in lateral view of embryos. **i** Quantitative comparison of the spine number per 2 mm^2^ between embryos injected with control and *fst* MO. Spine number in *fst* MO is lower than that in wild type. Abbreviations: BB, basibranchial; 1-5, ceratobranchials 1-5; CH, ceratohyal; M, Meckel’s cartilage. Scale bars, 100 μm (**c-f**), 50 μm (**g, h**).

### Signaling pathways perturbation during spine patterning and development

Spine initiation and development in pufferfish involves the expression of members of the Hh, BMP, Wnt and FGF signaling pathways (Fig. 3 and Supplementary Fig. 1). To understand the function of these pathways during spine development and also the potential role of Notch, an additional key developmental signaling pathway during spine development, we inhibited each pathway *in vivo* through the use of small molecules (pharmacological agents; e.g.^*56, 57*^). After hatching, embryos were exposed to small molecules in water for 72 hours, covering the key stages of early spine development (initiation, determination of patterning and initial primordium formation). Controls consisted of embryos treated with 1% DMSO. Following treatment, embryos were left to develop under standard conditions for a further 14 days. Embryos were then observed for phenotypic shifts following staining with Alcian Blue (Fig. 6a). Treatment with Hh antagonist, Cyclopamine (50 μM) severely repressed the formation of spines in the ventral region (Fig. 6b) - highlighting the need of Hh signaling for normal spine development. Next, treatment with DAPT (25 μM), an inhibitor of γ-secretase complex, which includes Notch pathway signaling, resulted in abnormal spine morphogenesis and a ca. 40% reduction in spine number in the ventral region of *Takifugu* embryos (Fig. 6c, l). This suggests that the Notch pathway is required for normal dermal spine formation. Given the result from the DAPT treatment, we then examined gene expression of Notch pathway members to determine whether Notch-related genes are expressed during spine development. We observed that at least one of the Notch ligands, *notch3*, is expressed in the specific territory where spines will form 10 dpf in wild-type embryos, although expression is not specifically restricted to the spine primordium (Fig. 6d). The same territory was also found to strongly express *shh*, which covers the skin surface in wild-type embryos at 10 dpf (Fig. 6e). Furthermore, we found that cyclopamine and DAPT treatments affect the activity of other pathways, such as the BMP and Wnt pathways 72 hours following the treatment period (Fig. 6f-k). In control embryos, *bmp2* and *lef1*, as markers for the pathways of BMP and Wnt, respectively, are expressed in an irregular pattern within the restricted ventral spine region (Fig. 6f, g), whereas following cyclopamine treatment, both genes showed greatly reduced expression patterns (Fig. 6h, i). Treatment with DAPT caused a reduction in *bmp2* expression, whilst *lef1* was significantly upregulated in the primordia (Fig. 6j, k). Treatment with the potent Bmp inhibitor LDN193189 (5 μM) and the FGF pathway inhibitor SU5402 (50 μM) lead to a modest reduction in the number of ventral spines in *Takifugu* (Fig. 6l).

**Fig. 6.**
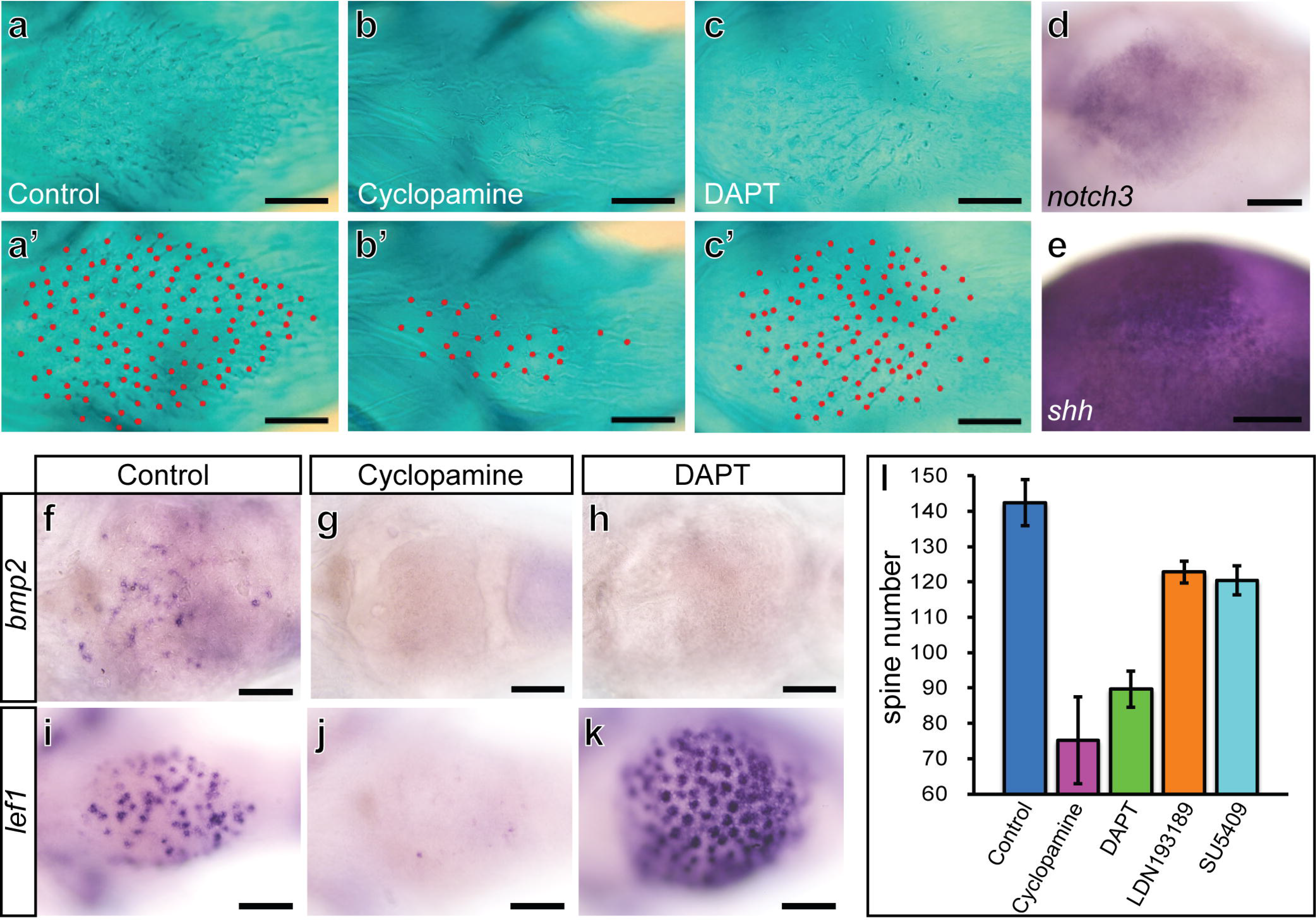
Manipulation of Hh, Notch, BMP and FGF pathways during spine development. **a-c** Alcian Blue stained ventral spine of *T. niphobles* embryos treated with small molecules for 72 hours and subsequent recovery process until 14 dpf. The treatment is 1% DMSO as a control (**a**), 50 μM Cyclopamine (**b**) and 25 μM DAPT (**c**) respectively. (**a’-c’**) Ventral spines in each sample marked with dotted red line. (**d, e**) Gene expression of *notch3* (**d**) and *shh* (**e**) in the ventral region where spines will form by whole mount RNA *in situ* hybridization of embryos at 10 dpf. (**f-k**) Comparison of gene expression in the spines for chemically treated embryos between 1% DMSO control (**f, g**), 50 μM Cyclopamine (**h, i**) and 25 μM DAPT (**j, k**) by whole mount RNA *in situ* hybridization of embryos past from 72 hours treatment. Gene expression for *bmp2* (**f, h, j**) and *lef1* (**g, i, k**). **l** Quantitative comparison for the spine total number of 14 dpf embryos between treatment with 1% DMSO control (n = 5), 50 μM Cyclopamine (n = 6), 25 μM DAPT (n = 9), 5 μM LDN193189 (n = 9) and 50μM SU5402 (n = 10). Scale bar, 100 μm.

Treatment with the Wnt-inhibitor, IWR-1-endo, mostly resulted in lethality of embryos (1 μM), however a single individual survived the treatment and displayed a severe reduction in the number of spines (Supplementary Table 2). Although this result hints at the potential effect of Wnt pathway inhibition on spine development, this result requires further experimentation to determine the extent of the role of Wnt signaling during ectodermal patterning in *Takifugu*. These results suggest that the 5 major signaling pathways, Hh, BMP, Wnt, FGF and Notch are involved during the development and patterning of dermal spines in pufferfish; the Hh pathway plays a prominent role in initiation and patterning of spines in *Takifugu*, and is critical for the maintenance or regulation of BMP and Wnt pathway molecules. Chemical perturbation experiments provide an important tool to determine the more generic roles of signaling pathways in organogenesis, and offer a focal point from which to design more specific tests of signaling molecule function.

### edar is not associated with spine development in Takifugu

EDAR (Ectodysplasin A receptor) is a cell surface receptor for ectodysplasin A (EDA), which plays an important role in the development of skin (ectodermal) appendages and scales in teleost fish^*11, 12, 16, 58*^. EDAR has been consistently observed in relation to development and patterning of dermal armor in teleost fishes. Notably, this gene is well characterized in relation to the evolution of stickleback armor patterning^*14, 19*^. To determine whether *edar* is involved in the development of the related, yet distinct, dermal armor of spines in pufferfishes, we examined *edar* gene expression from embryonic to juvenile stages of *Takifugu*. Given the role of ectodysplasin genes in scale development and patterning in teleost fishes, we hypothesized that *edar* would be expressed during dermal spine development in *Takifugu*. In embryonic stages, *edar* was expressed in the superficial epidermal cells scattered over the entire skin surface (Fig. 7a) in a similar expression pattern to *shh* (Fig. 7b). However, surprisingly, *edar* was not expressed in the dermal spine primordia in *Takifugu* (Fig. 7c). These data suggest that *shh* and *edar* are both expressed in skin cells in *Takifugu*, however unlike standard teleost scales, *edar* is not associated with the initiation of dermal spines. In ventral regions where spines initiate, *edar* was also expressed in skin cells, however its expression was absent from protruding epithelial cells overlying spine primordia (Fig. 7c). Expression of *edar* was not observed in the skin where spines were undergoing differentiation (Supplementary Fig. 2). At the later juvenile stages, expression of *edar* was present in the epidermis, especially in cells adjacent to each spine tip (Fig. 7d). These observations suggest that *edar* is not required for the initiation and site specification for spine primordia.

**Fig. 7.**
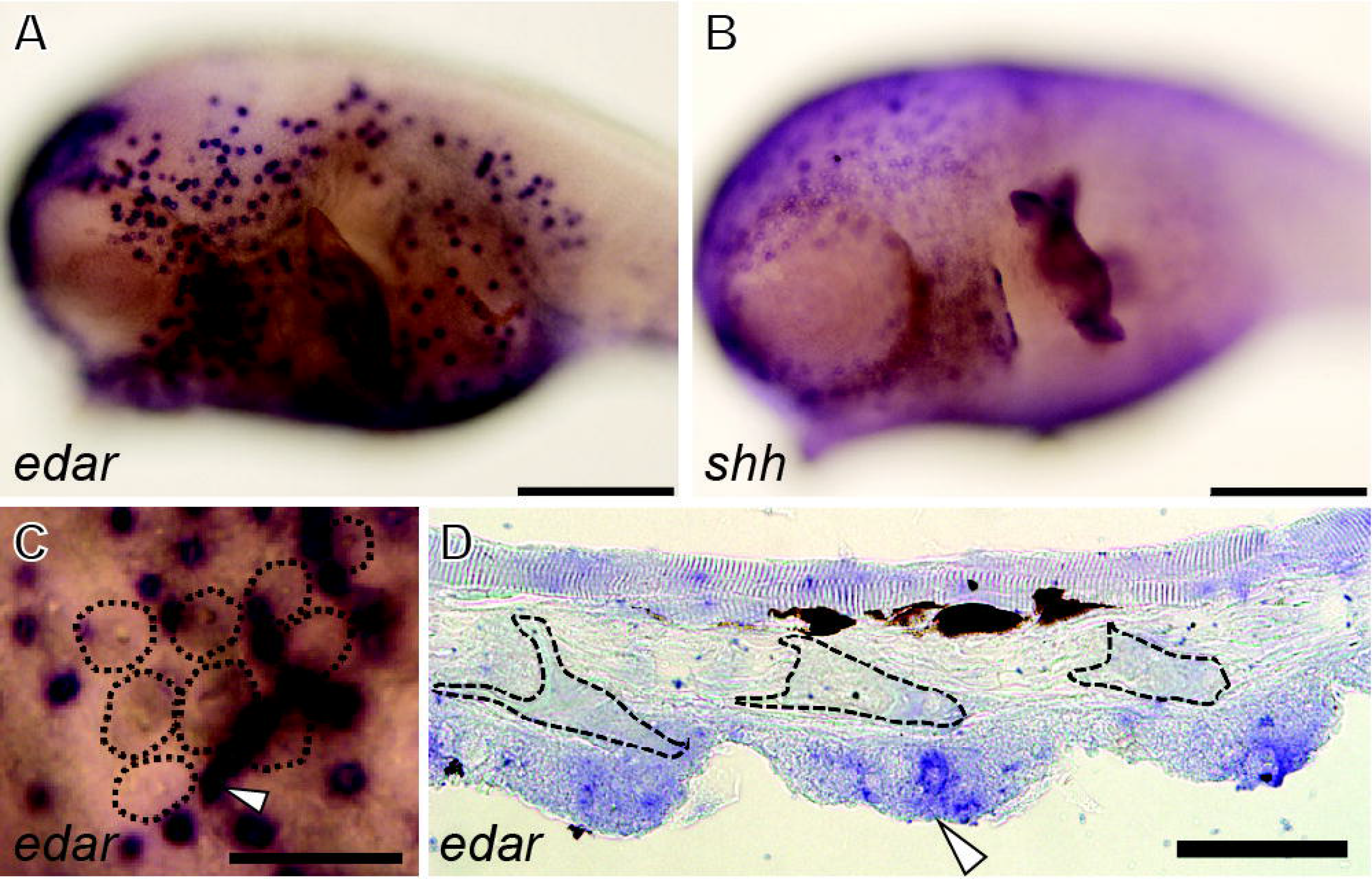
Gene expression patterns of *edar* and *shh* during spine development. *edar* (**a, c, d**) and *shh* (**b**). **a-b** Lateral view of 12 dpf embryo (**c**) Close-up of dorsal view of skin surface of a 12 dpf embryo. Black dotted lines indicate the area of individual spines. *edar* expressing cells are not associated with spines development (white arrowhead). (**d**) Transversal section in 50 dpf larva. Black dotted lines indicate the individual spine each. *edar* is expressed on the tip of the epithelium adjacent to spines (white arrowhead). Scale bars, 100 μM (**a, b**), 50 μM (**c, d**).

## Discussion

The pufferfish clade (Tetraodontidae) could have evolved a unique pattern of skin spines through changes to the initiatory process of skin appendage patterning, with early regional restriction of signaling molecules demarcating competent fields of spine formation. This is rather different from the typical signaling that instigates an initiatory row, from which many other appendage-types propagate, for example, in zebrafish scales, shark dermal denticles, reptilian scales and feather tracts. Spine primordia formation never start from a standard designation e.g. the midline of the trunk or in a caudal to rostral sequential pattern, instead pufferfish spine primordia form simultaneously within the territories established by the BMP, Wnt and Hh signaling molecules (Fig. 2). The earliest expression of Bmp, Notch and Wnt molecules seen in feather development, are not observed throughout the skin early in *Takifugu* development, however these molecules are only up-regulated in regions destined to become feather tracts and subsequent primordia*^52, 59^*. We observe molecules such as BMP, Notch and Hh genes expressed in a specific territory at the earliest stage of spine patterning that may set up the modified patterning of spines (Fig. 4, 6d-e). It is important to note that even within the pufferfish clade, spine coverage is not uniform, with species showing complete body coverage (e.g. *Carinotetraodon*) and others with a more reduced occurrence (e.g. *Takifugu*), therefore it appears that shifts in the competent territory for spine induction is flexible between species (Supplementary Fig. 4). Pufferfish offer a set of interesting developmental models in which to understand how regional restriction and elaboration of skin appendages has occurred during vertebrate evolution.

Morphological variation of the dermal skeletal ornament in Tetraodontiformes is substantial (Fig. 1). Triacanthodidae, the most basal family in the Tetraodontiformes phylogeny, possess numerous spines with complex morphology in the adult, however in the juvenile stages they exhibit a single spine that projects out from the cycloid compartment of the scale (Fig. 1a-c). This suggests that the common ancestor of this group had a spine attached to a scale with a cycloid base. This is similar in morphology to spinoid scales observed in other teleost groups, where spines project out from the scale posteriorly as continuations of the cycloid base^*23*^.

Based on this, we propose a hypothesis for the morphological evolution of the dermal skeleton in Tetraodontiformes (Fig. 8). Within this group, modifications to the ancestral scale have taken place, with notable diversification of the skin ornament. Different groups exhibit loss of either the cycloid base (i.e. Tetraodontidae and Diodontidae) or spines from the scale (i.e. Molidae and Ostraciidae). In molids and ostraciids, the cycloid base has been modified into a thick armor of polygonal plates, which may provide increased protection from predators. When looking at the diversification of spines, the Triacanthodidae, Balistidae, Monacanthidae and Triodontidae show a “cycloid” base adorned with a varying number of spines, whereas, the cycloid compartment has been lost in the Tetraodontidae and Diodontidae. We therefore suggest that spines in the Tetraodontidae and Diodontidae are highly derived dermal elements that are homologous to the standard scales of teleost fishes.

**Fig. 8.**
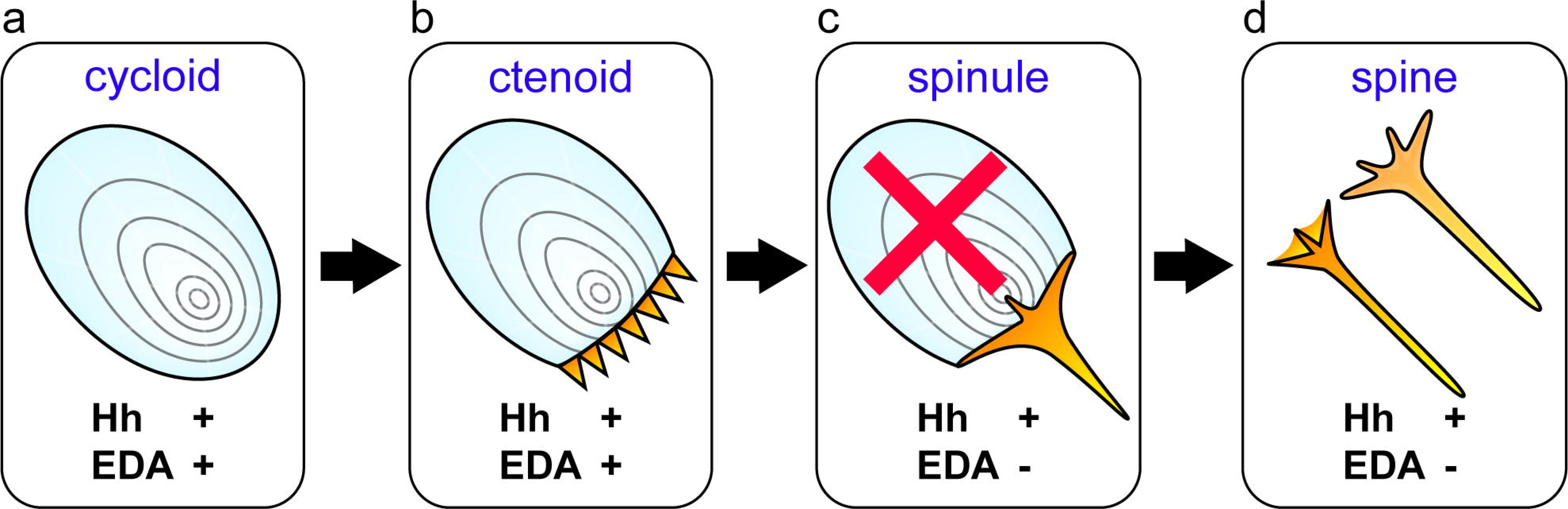
Schematic hypothesis of the Tetraodontidae and Diodontidae skin spines from scales. **a** Basic cycloid scale, of squamous type without ornamentations observed in teleost species. **b** Ctenoid scale with tooth-like spines projecting on the posterior side separate from the main cycloid compartment^*20*^. Cycloid and ctenoid scale require both Hedgehog (Hh) and EDA pathway for their development. **c** Plesiomorphic condition of tetraodontiform spinoid scale, composed of cycloid scale and single projecting spine posteriorly as in extant Triacanthodidae, Triacanthidae, Balistidae, Monacanthidae (see Fig. 1). During evolution of ancient lineage of Tetraodontidae and Diodontidae (or Molidae; Gymnodontes), cycloid compartment has been lost, but single spines are retained. **d** Modification of gene pathways, especially EDA, contributed to developmental changes forming scales without a “cycloid” scale compartment.

The EDA pathway could be a candidate set of molecules crucial to the diversification of tetraodontiform scales and appendage-types. The EDA pathway plays a role in fine-tuning the size, spacing, position and the shape of skin appendages in vertebrates^*60*^. For example, in mouse hair development, the absence of Eda (Eda-A1) results in aberration of primal hair, abnormal shape and decreased density, whereas overexpression of Eda leads to increased size and abnormal shape of hair^*61, 62*^ EDA signaling controls density and patterning of bony armor (modified scale) in several stickleback species^*19, 63*^, whilst mutation of *edar* in Medaka leads to significant reduction in the number of scales^12^. The EDA signaling pathway interacts with other signaling pathways such as Hh, Wnt, FGF and BMP in the development of skin appendage in many vertebrates^*14, 64–66*^. Therefore, a shift in EDA signaling is likely a key factor for morphological evolution of the dermal skeleton in Tetraodontiformes and other teleost groups.

## Conclusion

Scale morphology, especially in the highly derived superorder Acanthopterygii possesses various ornamentations on the posterior region of the cycloid scale, such as tubercles, ridges, serrations, cteni and spines^*67, 68*^. This implies that scales have evolved by acquisition of these ornamentations based on the original cycloid scale-type. Tetraodontiformes may be an exceptional example of scale evolution, especially with respect to the spines observed in Tetraodontidae and Diodontidae. We suggest these spines are formed through loss of the cycloid scale compartment present in the ancestor, to a reduced single spine, driven through drastic modification of gene pathway signaling during scale development (e.g. EDA signaling; Fig. 8). Tetraodontiform fishes have utilized this ‘plastic’ dermal system associated with BMP signaling to develop and modify spine and scale morphology to provide great variety and diversity of form. However, the key controlling factor(s) of this dermal diversification are yet to be determined. Taken together, these data suggest that EDA signaling may play a role in at least the scale to spine transition. In addition to the potential role of EDA, tinkering of the variety of signaling pathways at various stages of dermal armor develop (i.e. BMP signaling) has led to the diversity in patterning and unit morphology in this group of derived teleost fishes. *Takifugu* and *Tetraodon* pufferfishes have intriguing clues from their genomic sequences to suggest some mechanism for their diversity^*25*^, but this is not the whole story. Further studies on non-coding DNA, including transposable elements^*69*^ and genetic regulation will provide further clues toward understanding the evolution and diversity of form, seen to an extreme in this incredible assortment of fishes.

## Materials and Methods

### Animals

Eggs of the pufferfish *Takifugu niphobles* and juveniles of the filefish *Stephanolepis cirrhifer* were collected on Arai beach, Kanagawa prefecture, Japan. *Takifugu* eggs were raised to desired embryonic stages in fresh seawater at 20°C. Collections of *Diodon holocanthus* juveniles fixed with 10% formalin in seawater were supplied by Shimonoseki Marine Science Museum. Our comparative cleared and double stained methods were performed on tetraodontiform material obtained from the collection at the Natural History Museum London, and includes the species; *Hollardia sp*. and *Paratriacanthodes herrei* (Triacanthodidae), *Tripodichthys oxycephalus* and *Pseudotriacanthus strigilifer* (Triacanthidae), *Balistes vetula* and *B. capriscus* (Balistidae), *Stephanolepis sp*. and *S. hispidus* (Monacanthidae), *Ostracion sp*. and *O. trigonus* (Ostraciidae), *Triodon macropterus* (Triodontidae), *Takifugu niphobles* (Tetraodontidae), *Diodon holocanthu* (Diodontidae) and *Ranzania laevis* (Molidae). Species used in this study are listed in Supplementary Table 3.

### Histology and clearing and double staining

*Takifugu* embryos for histology staining were fixed in 4% paraformaldehyde in PBS (Phosphate Buffered Saline) at 4°C overnight and dehydrated in a graded series of methanol. For paraffin embedding, dehydrated samples were incubated 20 minutes with isopropanol and two times for 30 minutes with Histo-Clear. Leica RM2145 microtome was used to section paraffin embedded samples at 14 μm. Slides were stained with 50% haematoxylin for 10 minutes and Alcian Blue solution (0.01% Alcian Blue, 70% ethanol and acetic acid) for 30 minutes. Stained slides were washed with double distilled water and mounted with Fluoromount (SIGMA). Finally mounted slides were photographed by using a BX51 Olympus compound microscope equipped with an Olympus DP71 camera.

*Takifugu* embryos and juveniles for clearing and staining were breached with breaching solution (1% H_2_O_2_, 5% formamide, 0.5x SSC, 75 mM NaCl, and 7.5 mM sodium citrate, pH 7.0). For Alizarin Red staining juveniles were placed into 0.5% trypsin solution at room temperature for overnight. After protein digest, calcified tissues were stained using 0.01% Alizarin Red S in 0.5%K0H solution for 3 hours. Cartilages were stained with Alcian Blue solution for overnight, and spines were also observed in the cartilage stained prepations. Cleared and stained fishes were washed with 0.5%KOH then graded into 80% glycerin. Additional specimens illustrated in Fig. 1 were prepared according to a published protocol^*70*^.

### Scanning electron microscopy

*Takifugu* embryos were fixed in 4% paraformaldehyde in PBS at 4°C overnight and dehydrated in a graded series of methanol. The embryos were critical-point dried using liquid CO_2_, mounted on a sample holder, covered with gold, and viewed using a Philips XL-20 scanning electron microscope.

### Microinjections

Antisense morpholinos (MOs) were injected into *Takifugu* eggs with a micro needle using an injector NARISHIGE IM 300 (NARISHIGE). 0.8 mM MOs against *fst*(*follistatin*) (5’-TTCAGCATCCCAAACATGATGGAGC-3’) were injected into 1cell stage of *Takifugu* eggs.

### Molecular cloning and probe synthesis

Total RNA was extracted from whole bodies of *Takifugu* (7 dpf) in RNA later (Sigma) were transferred to TRIzol (Invitrogen) and washed with RNeasy cleanup kit (Qiagen). Single-stranded cDNA was synthesized from total RNA using a RETROscript Reverse Transcription Kit (Ambion) according to the manufacturer’s instructions. The sequence of genes was identified from the genome database available at the International Fugu Genome Consortium (http://www.fugu-sg.org/project/info.html). *Takifugu* cDNA clone of *β-catenin, lef1, wnt7a, shh, sostdc1, fgf3, fgf10a, bmp2, bmp4, notch3, fst* and *edar* were isolated by PCR using following forward and reverse primes designed from Takifugu rubripes genomic sequence: *β-catenin*, 5’-CCCTGAGGAAGATGATGTGGACAA-3’ and 5’-ACAGTTCTGGACCAGTCTCTGGCTG-3’; *lef1*, 5’-CAGTCCCAAATACCAGATTCATATC-3’ and 5’-TCTTCTTCTTTCCATAGTTGTCTCG-3’; *wnt7a*, 5’-CTTTGTCTGGGAATTGTCTATTTG-3’ and 5’-AGGTGTTGCATTTAACGTAGCAG-3’; *shh*, 5’-GAAGGCAAGATCACAAGAAACTC-3’ and 5’-ACGTTCCCACTTGATAGAGGAG-3’; *sostdc1*, 5’-TCTCTGGTCCTCCTCGTGTC-3’ and 5’-CGACTGGTTGTGGTGAGCCC-3’; *fgf3*, 5’-GTTGAATTTGTTGGATCCGGTTAG-3’ and 5’-TGACCTTCGTCTCTTAACTCTCTTG-3’; *fgf10a*, 5’-GTGTAGATGGACAGTGACACAAGG-3’ and 5’-TTCTTGTTGCGCCACTCCGCCGAG-3’; *bmp2*, 5’-TTAGAAGCTTTCACCATGAAGAGTC-3’ and 5’ - TTCATCCAGGTAGAGTAAGGAGATG-3’; *bmp4*, 5’-AGAACAACCATGCTAGTCTGATA-3’ and 5’-GTGGCAGTAATAGGCTTGATAACC-3’; *notch3*, 5’-TCTGACTACACTGGAAGCTATTGTG-3’ and 5’-TTGCAGTCAAAGTTGTCATAGAGAC-3’; *fst*, 5’-CTCTTGTTCATGTGGCTTTGTC-3’ and 5’-AGTCTGAGTCCTCATCTTCATCATC-3’; *edar*, 5’-GTACTCCAAAGGGAAGTACGAAATC-3’ and 5’-AAGATCTTTCTCCTCCGACTCTG-3’. The PCR clone in the pGEM-T-Easy Vector (Promega) was used as a template, and a digoxigenin (DIG)-labeled antisense RNA probe was synthesized by in vitro transcription with T7/SP6 RNA polymerase (Promega) and DIG RNA labeling mix (Roche).

### Whole mount and section in situ hybridization

All *Takifugu* embryos for *in situ* hybridization were fixed in 4% paraformaldehyde in 0.01 M PBS at 4°C overnight. Fixed samples were dehydrated with graded series of methanol. Whole mount *in situ* hybridization was performed according to a published protocol^*71*^. Section *in situ* hybridization was carried out on paraffin sections, they were de-paraffinised, rehydrated and superheated with 10 mM citric acid in diethylpyrocarbonate (DEPC) treated double distilled H_2_O (pH 6). Slides were incubated in hybridization buffer (50% formamide, 5x SSC (Saline Sodium Citrate), 500 mg/ml yeast tRNA, 50 mg/ml heparin, and 0. 01% Tween-20, pH 6.0) containing DIG-labeled antisense RNA probe at 61°C overnight. The hybridized slides were washed 1hour with 2xSSCT (Saline Sodium Citrate and 0.01% Tween-20, pH7.0) and three times for 30 minutes with 0.2xSSCT at 51°C. After incubation for 1 hour with blocking solution (2% Blocking Reagent (Roche) in MABT) at room temperature, slides were immunoreacted overnight at 4°C with an antiDIG Fab fragment conjugated to alkaline phosphate (Roche, 1:5000). After several washes for 6 hours with MABT (100 mM maleic acid, 150 mM NaCl, and 0.01% Tween 20, pH 7.5) and two washes with NTMT (100 mM NaCl, 50 mM MgCl_2_, 100 mM Tris-HCl (pH 9.5), and 0.01% Tween-20), samples were treated with BM purple AP Substrate precipitating solution (Roche). When satisfactory coloration was achieved, slides were washed with double distilled water and mounted with Fluoroshield with DAPI (SIGMA).

### Treatment with small molecules

Stock solutions were prepared for each chemical treatment experiment using Dimethyl Sulfoxide (DMSO) as a solvent. All treatment of small molecules were based on and adapted from^*53*^. 8 dpf *Takifugu* embryos were split into small molecules in sea water for 3 days. Final concentration of small molecules (Cyclopamine, DAPT, LDN193189, SU5402 and Iwr-1 (MedChem Express)) are shown in Supplementary Table 2. After the treatment, embryos for *in situ* hybridization were fixed and for clearing and staining were washed extensively with sea-water and were raised for 4 days prior to sacrifice and fixation. The number of ventral spines was counted under a BX51 Olympus compound microscope.

## Author Contributions

TS, DK, APT and GJF designed the research. TS, DK, APT, RB and GJF analysed the data. TS, DK, APT, RB and GJF performed the research. TS, AT, RB and GF wrote the paper. All authors edited and agreed on the final version.

## Acknowledgments

We thank members of the Fraser lab for comments on the manuscript and discussions, and Serina Hayes for laboratory assistance. We are grateful to Hiroyuki Doi, Toshiaki Ishibashi and Hisanori Kohtsuka for the donation of embryos. We also appreciate the open source CT scanning by members of the ‘Scan All Fishes’ consortium, available at Morphosource.org, for the *Takifugu oblongus* scan (used for Figure 2F), and especially Adam Summers and Matt Kolmann (University of Washington). This work was generously funded by the following research support: The Leverhulme Trust Research Project Grant RPG-211 (to G.J.F), Natural Environment Research Council (NERC) Standard Grant NE/ K014595/1 (to G.J.F), The Royal Society Research Grant RG120160 (to G.J.F), The Great Britain Sasakawa Foundation (to G.J.F and T.S) and the Daiwa Anglo-Japanese Foundation (to G.J.F). This work was also funded through “Adapting to the Challenges of a Changing Environment” (ACCE), an NERC funded doctoral training partnership (to A.T) ACCE DTP (NE/L002450/1).

## Supplementary Materials

**Supplementary Fig. 1**

Gene expression patterns during differentiation and development of spines. Transversal sections of hybridized spines in *Takifugu niphobles* embryos 15 days post fertilization illustrating the expression patterns of *β-catenin* (**a**), *lef1*(**b**), *wnt7a* (**c**), *shh* (**d**), *sostdc1* (**e**), *fgf3* (**f**), *fgf10a*(**g**), *bmp2*(**h**), *bmp4* (**i**) and *fst* (**j**). Black dotted lines indicate the individual spine each. **a-c** *β-catenin* and *wnt7a* are expressed in the epithelium uniformly surrounding spines, on the other hand, *lef1* is expressed in the posterior epithelia within the spine. **d** The expression of *shh* is more restricted within the epithelium compared to Wnt-related gene. **e** *sostdc1* is expressed in the epithelium anterior to the spine during morphogenesis. **f-g** *fgf3*and *fgf10a*, are extensively expressed in the epithelium adjacent apical tip of the spines. **h-j** *bmp2 and bmp4* are also extensively expressed the epithelium adjacent apical tip of spines, and *fst* is expressed within the epithelium around spines but not at the apical tip region. Scale bar, 25 μm.

**Supplementary Fig. 2**

Gene expression patterns of *edar* in 15 dpf *Takifugu* embryos. Whole mount RNA *in situ* hybridization of *edar* in the ventral (**a**) and lateral (**b**) region of a *Takifugu* embryo. *edar* is expressed in the epithelium adjacent apical tip of spines and neuromast but not cells entirely of the spines seen in 12 dpf embryos. Scale bar, 200 μm.

**Supplementary Fig. 3**

Schematic diagram representing gene expression during morphogenesis of spines. Summary of gene expression patterns by whole mount and section *in situ* hybridization observed in *Takifugu* embryos at 12 dpf (**a**) where spine primordia form (seen in Fig. 3) and 15 dpf (**b**) where spines develop during morphogenesis (seen in Supplementary Fig. 1).

**Supplementary Fig. 4**

Hypothesis of gene expression patterns in the skin that determines the spatial pre-pattern of spines. In the species of Diodontidae and Tetraodontidae, spine coverage is not uniform; **a** with species showing complete body coverage (e.g. *Diodon* and *Carinotetraodon*) and **b** others a more reduced occurrence (e.g. *Takifugu*). Molecules such as Hh, BMP, and Notch genes expressed in specific territories may set up the modified patterning of spines.

**Supplementary Table. 1**

List of scale structures amongst Tetraodontiformes. The scale typically consists of two regions, an anterior “cycloid” plate and posterior spinous structures. Adult scale morphology as observed in a published data^*18*^.

**Supplementary Table 2**

List of chemical perturbation experiments to inhibit signaling pathways during the development of spines. The phenotype in chemically treated embryos are reported by calculating average number of spines. Abbreviation; SD, Standard Deviation.

**Supplementary Table 3**

List of species used in this study for clearing and staining represented in Fig. 1. Institution abbreviation used in Table: Natural History Museum London (BMNH), Museum of Comparative Zoology, Harvard (MCZ), Muséum national d’histoire naturelle, Paris (MNHN) and Southeast Area Monitoring and Assessment Program Ichthyoplankton Archiving Center, Fish and Wildlife Research Institute (SEAMAP).

## References

1. Wu, P. et al. Evo-devo of amniote integuments and appendages. Int. J. Dev. Biol. 48, 249–270 (2004).

2. Sire, J. Y. & Huysseune, A. Formation of dermal skeletal and dental tissues in fish: a comparative and evolutionary approach. Biol. Rev. 78, 219–249 (2003).

3. Mikkola, M. L. & Millar, S. E. The mammary bud as a skin appendages: unique and shared aspects of development. J. Mammary‥ Gland. Biol. Neoplasia. 11, 187–203 (2006).

4. Di-Poi, N. & Milinkovitch, C. The anatomical placode in reptile scale morphogenesis indicates shared ancestry among skin appendages in amniotes. Sci. Adv. 2, 1–8 (2016).

5. Wildelitz, R. B., et al. Biology of feather morphogenesis: a testable model for evo-devo research. J. Exp. Zool. 298B, 109–122 (2003).

6. Chang, C. et al. Reptile scale paradigm: evo-devo, pattern formation and regeneration. Int. J. Dev. Biol. 53, 813–826 (2009).

7. Biggs, L. & Mikkola, M. L. Early inductive events in ectodermal appendage morphogenesis. Sem. Cell. Dev‥ Biol. 25, 11–21 (2014).

8. Mongera, A. & Nusselein-Volhard, C. Scales of fish arise from mesoderm. Curr. Biol. 23, R338–339 (2014).

9. Shimada, A. et al. Trunk exoskeleton in teleosts is mesodermal in origin. Nat. Commun. 4, 1639 (2013).

10. Sire, J. Y. & Akimenko, M. A. Scale development in fish: a review, with description of sonic hedgehog (shh) expression in the zebrafish (Danio rerio). Int. J. Dev‥ Biol. 48, 233–247 (2004).

11. Harris, M. P. et al. Zebrafish eda and edar mutants reveal conserved and ancestral roles of ectodyplasin signaling in vertebrates. PLoS. Genet. 4, e1000206 (2008).

12. Kondo, S. et al. The medaka rs-3 locus required for scale development encodes ectodysplasin-A receptor. Curr. Biol. 11, 1202–1206 (2001).

13. Rohner, N. et al. Dupilication of fgfr1 permits fgf signaling to serve as a target for selection during domestication. Curr. Biol. 19, 1642–1647 (2009).

14. Brown, N. M., Summets, B., Jones, F., Brady, S. D. & Kingsley, D.M. A recurrent regulatory change underliying altered expression and Wnt response of the stickleback armor plates gene EDA. eLife 4, e05290 (2015).

15. Cheng, J., Sedlazek, F., Altmuller, J. & Nolte, A. Ectodysplasin signaling genes and phenotypic evolution in sculpins (Cottus). Proc. R. Soc. 282, 20150746 (2015).

16. Iwasaki, M., Kuroda, J., Kawakami, K. & H, Wada. Epidermal regulation of bone morphogenesis through the development and regeneration of osteoblasts in the zebrafish scale. Dev. Biol. 437, 105–119 (2018).

17. Albertson, R. C., Kawasaki, C. K., Tetrault, R. E. & Powder, E. Kara. Genetic analyses in Lake Malawi cichlids identify new roles for Fgf signaling in scale shape variation. Commun. Biol. DOI: 10.1038/s42003-018-0060-4 (2018).

18. Aman, J. A., Fulbright, N. A. & Parichy, M. D. Wnt/β-catenin regulates an ancient signaling network during zebrafish scale development. bioRxiv. 293233; doi: https://doi.org/10.1101/293233 (2018).

19. Colosimo, P. F. et al. Widespread parallel evolution in sticklebacks by repeated fixation of ectodysplasin alleles. Science. 307, 1928–1933 (2005).

20. Santini, F. & Tyler, J. C. A phylogeny of the families of fossil and extant tetraodontiform fishes (Acanthomorpha, Tetraodontiformes), upper Cretaceous to Recent. Zool. J. Linn. Soc. 139, 565–617 (2003).

21. Tyler, J. C. Osteology, phylogeny, and higher classification of the fishes of the order Plectognathi (Tetraodontiformes). NOAA. Tech. Rep. NMFS. Circ. 434, 1–422 (1980).

22. Fraser, G. J., Britz, R., Hall, A., Johanson, Z. & Smith, M. M. Replacing the first-generation dentition in pufferfish with a unique beak. Proc. Natl. Acad. Sci. USA 109, 8179–8184 (2012).

23. Roberts, C. D. Comparative morphology of spined scales and their phylogenetic significance in the teleostei. Bull. Mar. Sci. 52, 60–113 (1993).

24. Williamson, W. C. Investigations into the structure and development of the scales and bones of fishes. Phil. Trans. R. Soc. Lond. 141, 643–702 (1851).

25. Venkatesh, B., Gilligan, P. & Brenner, S. Fugu: a compact vertebrate reference genome. FEBS. Let. 476, 3–7 (2000).

26. Jaillon, O. et al. Genome duplication in the teleost fish Tetraodon nigroviridis reveals the early vertebrate proto-karyotype. Nature 431, 946–957 (2004).

27. Matsuura, K. Taxonomy and systematics of tetraodontiform fishes: a review focusing primarily on progress in the period from 1980 to 2014. Ichthyol. Res. 62, 72–113 (2015).

28. Matsuura, K. The Living Marine Resources of the Western Central Atlantic. (Food and Agriculture Organization of the United Nations, Rome, 2002).

29. Britz, R., Ali, A., Philip, S., Kumar, K. & Raghavan, R. First record from the wild of Carinotetraodon imitator in Peninsular India (Teleostei: Tetraodontiformes: Tetraodontidae). Ichthyol. Explor. Freshw. 23, 105–109 (2012).

30. Katayama, E. & Matsuura, K. Fine Structures of Scales of Ocean Sunfishes (Actinopterygii, Tetraodontiformes, Molidae): Another Morphological Character Supporting Phylogenetic Relationships of the Molid Genera. Bull. Natl. Mus. Nat. Sci. Ser. A 42, 95–98 (2016).

31. Cleland, J. On the anatomy of the short sun-fish (Orthagoriscus mola). Nat. Hist. Rev‥ Lond. New. Ser. 2, 170–185 (1862).

32. Hertwig, O. Ueber das Hautskelet der Fische. 3: Das Hautskelet der Pediculati, der Discoboli, der Gattung Diana, der Centriscidae, einiger Gattungen aus der Familie der Triglidae und der Plectognathen. Morphol. Jhb. 7, 1–42 (1882).

33. Rosen, N. Studies on the plectognaths. 4. The integument. Ark. Zool. 8, 1–29 (1913).

34. Rosen, N. Studies on the plectognaths. 5. The skeleton. Ark. Zool. 10, 1–28 (1916).

35. Kaschkaroff, D. N. Materialien zur vergleichenden Morphologie der Fische. Vergleichendes Studium der Organisation von Plectognathi. Bull. Soc. Imp. Nat. Moscow 27, 263–370 (1914).

36. Gauldie, R.W. ‘Plywood’ structure and mineralization in the scale of the ocean sunfishes, Mola mola and M. ramsayi. Tissue. Cell24, 263–266 (1992).

37. Fujita, S. Studies on the life history and culture of common pufferfishes in Japan. Nagasaki. Pref. Res. Stn. Rep. 2, 1–121 (1962).

38. Guellec, D. L., Morvan-Dubois, G. & Sire, J. Y. Skin development in bony fish with particular emphasis on collagen deposition in the dermis of the zebrafish (Danio rerio). Int. J. Dev. Biol. 48, 217–231 (2004).

39. Johnson, R.L. & Tabin, C. J. Molecular models for vertebrate limb development. Cell 90, 979–990 (1997).

40. Graham, A. & Richardson, J. Developmental and evolutionary origin of the pharyngeal apparatus. Evo. Devo 3, 24 (2012).

41. Fraser, G. J., Cerny, R., Soukup, V, Bronner-Fraser, M. & Streelman, J. T. The Odontode Explosion: The origin of tooth-like structures in vertebrates. BioEssays 32, 808–817 (2010).

42. Grigoryan, T., Wend, P., Klaus, A. & Birchmeir, W. Deciphering the function of canonical Wnt signals in development and disease: conditional loss- and gain-of-function mutations of beta-catenin in mice. Genes. Dev. 22, 2308–2341 (2008).

43. Hovanes, K. et al. β-catenin sensitive isoforms of lymphoid enhancer factor 1 are selectively expressed in colon cancer. Nat. Genet. 28, 53–57 (2001).

44. Vogel, A., Rodriguez, C., Warnken, C. W. & Izpisua-belmonte, J. C. Dorsal cell fate specified by chick Lmx1 during vertebrate limb development. Nature 378, 716–720 (1995).

45. Vaijosalo, M. & Taipale, J. Hedgehog: Functions and mechanisms. Genes. Dev. 22, 2454–2472 (2008).

46. Ahn, Y., Sanderson, B. W., Klein, O. D. & Krumlauf, R. Inhibition of Wnt signaling by Wise (Sostdc1) and negative feedback from Shh controls tooth number and patterning. Development 137, 3221–3231 (2010).

47. Yanagita, M. et al. USAG-1, a bone morphogenetic protein antagonist abundantly expressed in the kidney. Biochem. Biophys. Res. Commun. 316, 490–500 (2004).

48. Neubuser, A., Peters, H., Balling, R. & Martin, G. R. Antagonistic interactions between FGF and BMP signaling pathways: a mechanism for positioning the sites of tooth formation. Cell 90, 247–255 (1997).

49. Shi, Y. & Massague, J. Mechanisms of TGF-beta signaling from cell membrane to the nucleus. Cell 113, 685–700 (2003).

50. Bauer, H. et al. Follistatin and Noggin are excluded from the zebrafish organizer. Dev Biol 204:488–507 (1998).

51. Thompson, T.B., Lerch, T. F., Cook, R. W., Woodruff, T. K. & Jardetzky, T. S. The structure of the Follistatin: Activin complex reveals antagonism of both type I and type II receptor binding. Dev. Cell. 9, 535–543. (2005).

52. Jiang, T.-X., Jung, H.-S., Wildelitz, R. B. & Chuong, C.-M. Self-organization of periodic patterns by dissociated feather mesenchymal cells and the regulation of size, number and spacing of primordial. Development 126, 4997–5009 (1999).

53. Plikus, M. V. Cyclic dermal BMP signaling regulates stem cell activation during hair regeneration. Nature 451, 340–344 (2008).

54. Mou, C., Jackson, B., Schneider, P., Overbeek, P. A. & Headon, D. J. Generation of the primary hair follicle pattern. Proc. Natl. Acad. Sci. USA 103, 9075–9080 (2006).

55. Dal-Pra, S., Furthauer, M., Van-Celst, J., Thisse, B. & Thisse, C. Noggin1 and Follistatin-like2 function redundantly to Chordin to antagonize BMP activity. Dev. Biol. 298, 514–526 (2006).

56. Fraser, G. J., Bloomquist, R. F. & Streelman, J. T. Common developmental pathways link tooth shape to regeneration. Dev. Biol. 377, 399–414 (2013).

57. Thiery, A. P. et al. Spatially restricted dental regeneration drives pufferfish beak development. Proc. Nat. Acad. Sci. USA 114, E4425–E4434 (2017).

58. Botchkarev, V. A. & Fessing, M. Y. Edar signaling in the control of hair follicle development. J. Investig. Dermatol. Symp. Proc. 10, 247–51 (2005).

59. Pan, Y. Gamma-secretase functions through Notch signaling to maintain skin appendages but is not required for their patterning or initial morphogenesis. Dev. Cell. 7, 731–743 (2004).

60. Sadier, A., Viriot, L., Pantalacci, S. & Laudet, V. The ectodyplasin pathway: from diseases to adaptations. Trends. Genet. 30, 24–31 (2014).

61. Cui, C. et al. Inducible mEDA-A1 transgene mediates sebaceous gland hyperplasia and differential formation of two types of mouse hair follicles. Hum. Mol. Gen. 12, 2931–2940 (2003).

62. Mustonen, T. et al. Ecdysplasin A1 promotes placodal cell fate during early morphogenesis of ectodermal appendages. Development 131, 4907–4919 (2004).

63. Cresko, W. A. et al. Parallel genetic basis of repeated evolution of armor loss in Alaskan threespine stickleback populations. Proc. Natl. Acad. Sci. USA 101, 6050–6055 (2008).

64. Mikkola, M. L. et al. TNF superfamily in skin appendage development. Cytokine. Growth. Factor. Rev. 19, 219–230 (2008).

65. Haara, O. et al. Ecdysplasin regulates activator-inhibitor balance in murin tooth development through fgf20 signaling. Development 139, 3189–3199 (2012).

66. Haara, O. et al. Ectodysplasin and Wnt pathways are required for salivary gland branching morphogenesis. Development 138, 2681–2691 (2011).

67. Agassiz, L. Recherches sur les Poissons Fossiles. Neuchate1: XLIV. (1833–1844).

68. Sire, J. Y. Ontogenic development of surface ornamentation in the scales of Hemichromis bimaculatus (Cichlidae). J. Fish. Biol. 28, 713–724 (1986).

69. Volff, J. N., Bouneau, L., Ozouf-Costaz, C. & Fischer, C. Diversity of retrotransposable elements in compact pufferfish genomes. Trends. Genet. 19, 674–678 (2003).

70. Taylor, W. R. & van Dyke, G. C. Revised procedures for staining and clearing small fishes and other vertebrates for bone and cartilage study. Cybium 9, 107–119 (1985).

71. Shono, T., Kurokawa, D., Miyake, T. & Okabe, M. Acquisition of glial cells missing 2 enhancers contributes to a diversity of ionocytes in zebrafish. PLoS. ONE6, e23746 (2011).

